# KRAS Inhibition Activates an Actionable CD24 “Don’t Eat Me” Signal in Pancreatic Cancer

**DOI:** 10.1101/2023.09.21.558891

**Authors:** Yongkun Wei, Minghui Liu, Er-Yen Yen, Jun Yao, Zhenzhen Xun, Phuoc T. Nguyen, Xiaofei Wang, Zecheng Yang, Abdelrahman Yousef, Dean Pan, Yanqing Jin, Ching-Fei Li, Madelaine S. Theardy, Jangho Park, Yiming Cai, Mitsunobu Takeda, Matthew Vasquez, Elizabeth M. Park, David H Peng, Yong Zhou, Hong Zhao, Timothy P Heffernan, Andrea Viale, Huamin Wang, Stephanie S. Watowich, Han Liang, Dan Zhao, Ronald A. DePinho, Wantong Yao, Haoqiang Ying

## Abstract

KRAS^G12C^ inhibitors (G12Ci) have produced encouraging, albeit modest and transient, clinical benefit in pancreatic ductal adenocarcinoma (PDAC). Identifying and targeting resistance mechanisms to G12Ci treatment is therefore crucial. To better understand the function of KRAS^G12C^ and possible G12Ci bypass mechanisms, we developed an autochthonous KRAS^G12C^- driven PDAC model. Compared to the classical KRAS^G12D^ PDAC model, the G12C model exhibits slower tumor growth, yet similar histopathological and molecular features. Aligned with clinical experience, G12Ci treatment of KRAS^G12C^ tumors produced modest impact despite stimulating a ‘hot’ tumor immune microenvironment. Immunoprofiling revealed that CD24, a ‘don’t eat me’ signal, is significantly upregulated on cancer cells upon G12Ci treatment. Blocking CD24 enhanced macrophage phagocytosis of cancer cells and significantly sensitized tumors to G12Ci treatment. Similar findings were observed in KRAS^G12D^-driven PDAC. Together, this study reveals common and distinct oncogenic *KRAS* allele-specific biology and identifies a clinically actionable adaptive mechanism that may improve the efficacy of oncogenic KRAS inhibitor therapy in PDAC.

**Significance:** Generation of an autochthonous KRAS^G12C^-driven pancreatic cancer model enabled elucidation of specific effects of KRAS^G12C^ during tumor development, revealing CD24 as an actionable adaptive mechanism in cancer cells induced upon KRAS^G12C^ inhibition.

## Introduction

KRAS is the most frequently mutated oncogene in human cancer with hot-spot mutations at Glycine^12^, Glycine^13^ and Glutamine^61^ (1). While all oncogenic KRAS mutations result in the constitutive activation of KRAS protein, different alleles exhibit somewhat distinctive biochemical properties and tissue-specific distribution patterns (2,3). Oncogenic mutations in KRAS are present in the vast majority of pancreatic ductal adenocarcinoma (PDAC) with G12D (41%), G12V (34%) and G12R (16%) being the most frequent mutations, along with other less common mutations such as Q61H (4%) and G12C (1.5%) (4,5). Retrospective analysis suggests that, among the KRAS mutations commonly found in PDAC, the G12D mutation is associated with worst prognosis (6) which mirrors survival data in lung and colon cancer patients (2), suggesting allele-specific differences in the tumor biological impact of various oncogenic KRAS alleles. It should be noted that the preclinical studies used to establish the critical function of KRAS oncogene for PDAC initiation and maintenance have been based largely on models driven by either Kras^G12D^ or Kras^G12V^ (7–12). Together, these data emphasize the need for the study of oncogenic KRAS allele-specific biology in PDAC and other cancer types.

KRAS, long considered an undruggable target, has been targeted by drugs including a KRAS^G12C^ inhibitor (G12Ci) which is approved for the treatment of KRAS^G12C^-mutant non-small cell lung cancer (NSCLC) (13,14). In a phase II trial for KRAS^G12C^-mutated PDAC, AMG 510 (Sotorasib) showed promising anticancer activity, with a 21% response rate and over 70% patients showing some tumor shrinkage (15). Recently, a novel KRAS^G12D^ inhibitor, MTXT1133, showed strong efficacy in pre-clinical animal models (16,17). Although various KRAS inhibitors have transformed the treatment landscape of KRAS-driven cancers, the rapid re-emergence of tumors has severely limited their clinical benefit (18,19). Multiple studies have elucidated potential resistance mechanisms to G12Ci in NSCLC including accumulation of additional mutations in the KRAS^G12C^ oncogene, re-activation of the KRAS signaling through the hyperactivation of multiple receptor tyrosine kinases (RTKs), and cell state transformation through epithelial to mesenchymal transition (EMT) (14,20–26). However, the molecular mechanisms underlying the resistance to G12Ci in PDAC remain poorly understood, owing in part to the lack of relevant KRAS^G12C^-driven preclinical models. With respect to PDAC models, the KRAS^G12D^ model has proven capable of anticipating resistance mechanisms involving cancer cell intrinsic and paracrine mechanisms (27–29). Here, we have engineered a G12C PDAC model revealing a less aggressive tumor phenotype relative to the KRAS^G12D^-driven model and a novel bypass mechanism in the face of AMG 510 treatment. Co-targeting CD24, a ‘do not eat me’ signal (30) upregulated in AMG 510-treated cancer cells, sensitizes PDAC to KRAS inhibitors, informing a highly effective co-extinction hypothesis for PDAC patients.

## Materials and Methods

### Animal studies

All animal studies were carried out in accordance with and approved by the Animal Care and Use Committees at The University of Texas MD Anderson Cancer Center (IACUC protocol number: 00001549-RN02).

### Mice

P48Cre, LSL-Kras^G12D^, LSL-Kras^G12C^, and p53^L^ mice have been described previously (10,31–33). Mice were interbred and maintained on C57BL/6 background in pathogen-free conditions. The C57BL/6 mice were purchased from Jackson Lab (Jackson Laboratory, RRID: IMSR_JAX:000664). NCr Nude mice were purchased from Taconic Biosciences.

Murine PDAC cell lines were derived from tumor-bearing K^C^PC (49725, 50760) or K^D^PC (55582) mice on a congenic C57BL/6 background. 49725(1 × 10^6^) and 50760 (5 × 10^5^) or 55582 (3 × 10^5^) cells were suspended in 100 μL medium and injected subcutaneously into the right flank of C57BL/6 mice. Human PDAC cell line AsPC1 (RRID: CVCL_0152) was obtained from the American Type Culture Collection (ATCC) in 2023, maintained in RPMI supplemented with 10% FBS, and incubated at 37°C with 5% CO_2_. Cell line identity was verified by short tandem repeat (STR) analysis in 2023. All cell lines were regularly tested for mycoplasma using Mycolor One-Step Mycoplasma Detector (Vazyme) and found negative, and early passages (3–5) were used for experiments. Human PDAC cell line AsPC1 with KRAS G12D mutation (1 × 10^6^) were injected into nude mice using the same method. Tumor size was measured every 2 days using a caliper and calculated using the formula length × width^2^/2. When tumor size reached 20 to 80 mm^3^, mice were randomly grouped and administered treatment. For AMG 510 treatment, mice were administered 30 mg/kg AMG 510 [dissolved in 0.2% Hydroxypropyl methylcellulose + 0.1% Tween 80] by oral gavage daily until the tumor size reached limit. For CD24 antibody treatment, mice were injected intraperitoneally with 100 μg of CD24 antibody (clone M1/69; #BE0360; Bio X Cell) or control rat IgG (clone LTF-2; #BE0090; Bio X Cell) twice a week. For MRTX1133 treatment, mice were injected intraperitoneally with 30 mg/kg MRTX1133(dissolved in 10% Captisol in 50 mmol/L citrate buffer pH 5.0) daily. For nude mice CD24 antibody treatment, mice were injected intraperitoneally with 100 μg of CD24 antibody (ATG-031; cat# HY-P99176; MCE) twice a week. On day 21, tumor tissue (*n* = 3/group) was harvested and prepared for single-cell suspension for further analysis. The remaining mice were monitored until the tumor size reached the limit, then euthanized. Waterfall plots were calculated using the baseline tumor volume on day 0 compared with tumor volume at the end of the experiment. The percent change in tumor volume was calculated by ((final volume – initial volume)/initial volume) × 100.

### Macrophage Depletion

Once tumors reached an average volume of 20 to 80 mm^3^, AMG 510 (30 mg/kg) and anti-CD24 were administered for 2 weeks. Macrophage depletion started 1 day before initiating AMG 510 and anti-CD24 treatment and continued every other day for the remainder of the experiment. Depleted mice received 400 μg of CSF1R (Clone AFS98) antibodies i.p., whereas control mice were given isotype control (MOPC-21; BE0083; Bio X Cell). Peripheral blood and tumor tissues were collected to confirm macrophage depletion via flow cytometry.

### Immunoblots

Cells were lysed in RIPA buffer (50mM Tris-HCL, pH 7.6, 150mM NaCl, 1% NP-40, 0.5% Sodium deoxycholate, 0.1% SDS, 1mM EDTA) supplemented with protease inhibitors (Complete Mini, Roche). To prepare the whole cell lysates, 6 × SDS sample buffer was directly added to the cell lysates before resolved on SDS-PAGE and immunoblotted with primary antibodies. The protein concentrations of the lysates were measured using the Bio-Rad protein assay reagent. After washing, the membrane was incubated with horseradish peroxidase– conjugated secondary antibody. After washing, the membrane was developed with enhanced chemiluminescence. Ras-GTP levels were assessed by Active Ras Pull-Down and Detection Kit using RAF-RBD fused to GST to bind active (GTP-bound) RAS. Protein lysates were incubated with glutathione resin and GST protein-binding domains to capture active small GTPases according to the manufacturer’s protocol. After washing, the bound GTPase was recovered by eluting the GST-fusion protein from the glutathione resin. The purified GTPase was detected by Western blot analysis using mouse monoclonal anti-KRAS antibody.

### Real-time Quantitative PCR

Total RNA was isolated using RNeasy Mini Kit (Qiagen), and cDNA was synthesized using SuperScript™ III First-Strand Synthesis Super Mix (Thermo Fisher Scientific). qPCR reactions were performed using Fast SYBR Green Master Mix (Thermo Fisher Scientific) on BIO-RAD CFX96 Real-Time PCR system. The threshold cycle (Ct) value for each gene was normalized to the expression levels of ACTIN, and relative expression was calculated by normalizing the delta Ct value. The primer sequences (5’–3’) for the transcripts analyzed are as follows: *CD24* forward: CTTCTGGCACTGCTCCTACC *CD24* reverse: GAGAGAGAGCCAGGAGACCA *ACTIN* forward: GGCTGTATTCCCCTCCATCG *ACTIN* reverse: CCAGTTGGTAACAATGCCATGT

### Cell Viability Assay

To determine IC_50_ values, 1 to 3 × 10^3^ cancer cells were plated in a 96-well plate. Cells were treated 24 hours later with DMSO or serial dilutions of AMG 510 or MRTX1133. Cell viability was read 72 hours later using MTT. IC_50_ values were generated using GraphPad Prism (RRID:SCR_002798) version 9.1.2.

### CyTOF analysis

Mouse Tumor Dissociation Kit (130-096-730, Miltenyi Biotec) and gentleMACSOcto Dissociator (130-096-427, Miltenyi Biotec) were used to digest the excised tumors from mice. Cells were then blocked with CD16/CD32 (156604; Biolegend) antibody followed by incubation with a mixture of metal-labeled antibodies (Supplementary Table S1) for 1 hour at room temperature. Cisplatin (195Pt, Fluidigm) was used as a marker to detect dead cells. Cell-ID Ir-intercalator (Fluidigm) was incubated overnight at 4°C. The samples were analyzed by using the Helios System (Fluidigm) from MD Anderson Flow Cytometry and Cellular Imaging Core Facility. Data was processed by the FlowJo (RRID:SCR_008520) software and Cytobank (RRID:SCR_014043; Cytobank, Inc.).

### IHC staining

Murine tumors were fixed overnight in 10% formalin. After 24 hours, tissues were transferred to 70% ethanol and processed for paraffin embedding. Tissues were sectioned at 5 μm. For immunofluorescence staining, slides were deparaffinized and rehydrated through a series of xylene and ethanol washes. Slides underwent antigen retrieval (Antigen Retrieval Citra Plus solution, Biogenex #HK080, 95 °C for 15 min) followed by blocking with 5% goat serum and 1% BSA in PBS 0.1% Triton-X100 for 1 hour at room temperature. Primary antibodies were added to the slides and incubated overnight at 4°C. Slides were then washed 2× with PBS 0.1% Triton-X100 and incubated with secondary antibodies (Alexa Fluor secondaries, 1:150) for 1 hour at room temperature. Nuclei were counterstained with DAPI (1:1,000). Slides were mounted with ProLong Diamond Antifade Mountant and coverslipped. Images were scanned using PE Vectra Polaris. QuPath and ImageJ (RRID:SCR_003070) software was used for quantitation. For IHC staining, antigen retrieval was carried out by heating for 12 minutes in DAKO Target Retrieval Solution using microwave. Sections were incubated with 3% H_2_O_2_ for 30 min at room temperature to block endogenous peroxidase activity. After incubating in 10% normal goat or horse serum for 1 h to block non-specific binding of IgG, sections were incubated with primary antibody at 4°C overnight. After wash with PBS, sections were then incubated for 1 h with ImmPRESS-HRP polymer reagent (Vector). Samples were developed with DAB quanto chromogen substrate (Epredia) and counterstained by hematoxylin. To measure protein expression by IHC, stained sections were analyzed at ×400 resolution and scored by H-score system. An H-score (range, 0– 300) was calculated as the sum of the highest intensity of staining (0, negative; 1, weak positive; 2, moderate positive; 3, strong positive) and percentage of positive cells (0%–100%; with any intensity of positive staining).

### Flow Cytometry Analysis

In vitro cultured cells or single-cell suspension from mouse tumors were stained for flow cytometry analysis. Mouse Tumor Dissociation Kit (130-096-730, Miltenyi Biotec) and gentleMACSOcto Dissociator (130-096-427, Miltenyi Biotec) were used to digest the excised tumors from mice. Samples were then washed with DMEM and 10% FBS and filtered through a 70-μm strainer to obtain a single-cell suspension. Cells from mouse tissues were blocked with CD16/CD32 (156604, Biolegend). Cells were labeled with primary fluorophore-conjugated antibodies for 1 h at 4°C. Cells were washed and resuspended in flow buffer. For intracellular staining, cells were permeabilized and fixed for 30 minutes at 4°C, washed, and then stained for intracellular markers for 1 h at 4°C. Data were acquired on a BD FACSCanto II flow cytometer, with analysis performed using FlowJo (RRID:SCR_008520) version 10.9.0.

### GeoMx digital spatial profiler (DSP) for RNA profiling in FFPE tissues

Five-µm FFPE tissue sections from 4 mice (2 G12C and 2 G12D) were mounted on positively charged histology slides. Slides were deparaffinized, rehydrated and incubated for 15 min in 1× Tris-EDTA pH 9.0 buffer at 100 °C. Slides were then incubated in 1 µg ml^−1^ proteinase K (Thermo Fisher Scientific, AM2546) in PBS for 15 min at 37 °C. Tissues were then fixed in 10% neutral-buffered formalin for 5 min, incubated in NBF stop buffer (0.1 M Tris Base, 0.1 M Glycine, Sigma) for 5 min twice, then washed in PBS. Tissues were then incubated overnight at 37 °C with GeoMx RNA detection probes in Buffer R (Nanostring Technologies). After washing, tissues were blocked in Buffer W (Nanostring Technologies) for 30 min at room temperature. Next, antibodies targeting PanCK and CD45 (Nanostring Technologies) in Buffer W were applied to each section for 1 h at room temperature. Slides were washed twice in fresh 2× SSC then loaded on the GeoMx Digital Spatial Profiler.

In the process entire slides were imaged at ×20 magnification and a total of 73 circular ROIs with a 200 μm diameter were selected and the GeoMx software was used to define AOIs (or segments) as one segment containing positive immunofluorescent signal for PanCK.

Once AOIs were defined, the DSP then exposed ROIs to 385 nm light (UV), releasing the indexing oligonucleotides, which were collected with a microcapillary. Indexing oligonucleotides were then deposited in a 96-well plate for subsequent processing.

Sequencing libraries were generated by PCR from the photo-released indexing oligos and AOI-specific Illumina adapter sequences, and unique i5 and i7 sample indices were added. PCR reactions were pooled and purified using AMPure XP beads (Beckman Coulter, A63881), according to the manufacturer’s protocol. Pooled libraries were sequenced at 2 × 75 base pairs and with the single-index workflow on an Illumina NextSeq to generate 458 M raw reads.

PCA analysis and unsupervised clustering analysis were performed using R software (prcomp function and pheatmap package). Differential gene expression analysis was carried out using R DESeq2 package with parameters of fold change > 1.5 and adjusted p value < 0.05. GSEA analysis was performed using R fgsea package.

### In vitro and in vivo phagocytosis assay

Flow cytometry-based phagocytosis assay was performed according to literature (30).

*In vitro* phagocytosis assays were performed by co-culture target cells and macrophages at a ratio of 100,000 target cells to 50,000 macrophages for 2 h in a humidified, 5% CO_2_ incubator at 37°C in ultra-low-attachment 96-well U-bottom plates (Corning) in serum-free medium. Cells with endogenous GFP were harvested from plates using TrypLE Express (Life Technologies) prior to co-culture. Macrophages (RAW264.7 mouse macrophage cell line) (RRID: CVCL_0493) were harvested from plates using TrypLE Express. Monoclonal antibody CD24 (Clone M1/69, BioXcell) and isotype controls were added at a concentration of 10 μg/mL. After co-culture, phagocytosis assays were stopped by placing plates on ice, centrifuged at 400 g for 5 min at 4°C and stained with PE-labeled anti-F4/80 to identify mouse macrophages. Assays were analyzed by flow cytometry. Phagocytosis was measured as the number of F4/80+, GFP+ macrophages, quantified as a percentage of the total F4/80+ macrophages. Each phagocytosis reaction was performed in a minimum of technical triplicate.

For In vivo phagocytosis assay, female C57BL/6 mice, 6–8 weeks of age, were injected with 5 × 10^5^ 50760-GFP cells into the right flank. Tumors were treated with vehicle plus IgG or AMG 510 plus anti-CD24 antibody for 14 days, after which tumors were resected and dissociated mechanically and enzymatically as described above. Single-cell suspensions of tumors were blocked using anti-CD16/32 (156604, BioLegend) for 15 min on ice. Phagocytosis was measured as the percentage of PerCp-F4/80^+^ macrophages that were also GFP^+^.

### Multiplex immunofluorescence staining

FFPE slides were deparaffinized in xylene and rehydrated in an alcohol gradient. Tissue underwent citrate-based antigen retrieval (Antigen Retrieval Citra Plus solution, Biogenex #HK080, 95 °C for 15 min) before blocking with 5% goat serum and 1% BSA in PBS 0.1% Triton-X100 for 1 h. Slides were incubated with unconjugated primary antibodies for 1 h at room temperature, followed by conjugated secondary antibodies for 1 h at room temperature, then with conjugated primary antibodies overnight at 4 °C. Slides were washed with PBS 0.1% Triton-X100 between each step. Nuclei were counterstained with DAPI (1:1,000). Slides were mounted with ProLong Diamond Antifade Mountant and cover slipped. Images were scanned using PE Vectra Polaris. Slides were incubated with stripping buffer (62.5 mM Tris Base / Tris-HCl, 2% Sodium Dodecyl Sulfate, 0.8% Beta ME, 55°C, 30 min) and then stained with another round of antibodies. There were 4 rounds of staining and slides scan. The antibodies used were listed at Supplementary Table S1. Image registration was conducted using MATLAB (RRID:SCR_001622) (The MathWorks, Inc.). Initially, the DAPI image from each imaging cycle was utilized to compute a transformation matrix using SURF features (https://doi.org/10.1016/j.cviu.2007.09.014). Subsequently, the extracted features were matched, employing the following parameters: Metric Threshold = 505, Num Octaves = 4, Num Scale Levels = 6. Automated image segmentation and quantification were performed using MATLAB (The MathWorks, Inc.). Specifically, Otsu’s thresholding method and the marker-controlled watershed algorithm were applied to segment and define the nuclear regions of individual cells. Subsequently, the expression of each marker within the segmented areas was quantified based on pixel intensity. Single-cell marker expression data underwent clustering using the R implementation of Phenograph (cytofkit, https://doi.org/10.1371/journal.pcbi.1005112, 10.1088/1742-5468/2008/10/P10008) with the parameter k=15. Prior to clustering, CD45^+^ Amylase^-^ cells in the lesion sites are selected, and marker expression values were normalized to the 99th percentile across all cells for each channel independently. Following clustering, the resulting clusters were labeled based on their marker expression signatures.

### RNA-seq analysis

Total RNA was isolated using RNeasy Mini Kit (Qiagen). The library preparation, sample clustering and sequencing were done in MD Anderson ATGC core. The sequencing was carried out on the Illumina NovaSeq platform. Differential expression analysis of two groups in RNA-seq was performed using the DESeq2 R package. The resulting *P* value was adjusted using the Benjamini and Hochberg approach for controlling the false discovery rate. Genes with an adjusted *P* < 0.05 found by DESeq2 were assigned as differentially expressed. Gene function enrichment analysis was done with clusterProfiler (RRID:SCR_016884) by R.

### Single cell RNA sequencing analysis

Tumors were dissected from mice and single-cell suspensions were obtained. Cells were counted and loaded onto the 10X device (10X Genomics). Samples were processed per the manufacturer’s protocol and sequenced on an Illumina NextSeq sequencer. Single cell RNAseq raw data (fastq files) were mapped to mouse transcriptome mm10 using cellranger v7.1.0. One sample with mapped data to the same reference genome was downloaded from NCBI GEO (https://www.ncbi.nlm.nih.gov/geo/, GSE125588, sample GSM3577885). Samples were processed using R SEURAT (RRID:SCR_007322) package to filter off poor quality cells based on mitochondria content and total gene numbers. Cells from all samples were merged and integrated by Seurat to remove batch effects. Cells were then clustered and annotated using cell markers from Zhang et al (34).

Differential gene expression analysis was performed using R DESeq2 (RRID:SCR_000154) package with p value cutoff of 0.05 and fold change cutoff of 1.5. GSEA analysis was performed for activated pathways using R fgsea package using ontology annotations from MSigDB (https://www.gsea-msigdb.org/gsea/msigdb) with a p value cutoff of 2e-3.

### Statistical analysis

GraphPad Prism (RRID:SCR_002798) version 9.1.2 was used for statistical analyses. Data are presented as means ± SD or SEM. Two-tailed Student *t* test and two-way ANOVA with multiple corrections were performed for comparison between groups. For IC_50_ generation, concentrations were log transformed, data were then normalized to control, and log(inhibitor) versus response (four parameters) test was used. A *P* value < 0.05 was considered statistically significant.

### Data availability

The RNA-seq data generated in this study are publicly available in Gene Expression Omnibus (GEO) at GSE303279 and GSE307328. The scRNA-seq data generated in this study are publicly available in Gene Expression Omnibus (GEO) at GSE305316 and GSE307209. The previously published scRNA-seq data analyzed in this study were obtained from Gene Expression Omnibus (GEO) at accession number GSE125588, sample GSM3577885. All other raw data generated in this study are available upon request from the corresponding author.

## Results

Pancreatic-specific expression of *Kras^G12D^* from the endogenous allele results in the development of premalignant lesions, such as PanINs and acinar-to-ductal metaplasia (ADM), with the occasional progression to invasive carcinoma at low penetrance after a long latency period (10). Deletion or mutation of additional tumor repressors such as *Ink4a* or *Trp53* cooperates with *Kras* oncogene to accelerate the tumor progression and leads to the development of invasive PDAC at full penetrance (7,11,35). Knock-in of Kras^G12C^ into mouse pancreas also induces the formation of premalignant lesions (31). To ensure full malignant progression and assess the role of Kras^G12C^ in advanced PDAC and therapy response, we established a mouse model engineered with the *LSL- Kras^G12C^*, conditional *p53* knockout (*p53^L^*), and pancreatic-specific *p48-Cre* alleles (31–33). The resulting *p48-Cre LSL-Kras^G12C^ p53^L/+^* (K^C^PC) mice exhibit decreased number and area of PanIN/ADM lesions compared to age-matched Kras^G12D^-driven controls (*p48-Cre LSL-Kras^G12D^ p53^L/+^*, K^D^PC) at 8, 12 and 16 weeks (Fig.1A-C). The decreased neoplastic activity of *Kras^G12C^* aligns with decreased downstream signaling (phospho-ERK and phospho-AKT) and cell proliferation (Ki67) in premalignant epithelial cells (Fig.1D-G; Supplementary Fig. S1A, B), suggesting that KRAS^G12C^ is a less potent oncogenic allele compared to KRAS^G12D^ for PDAC development.

**Figure 1.**
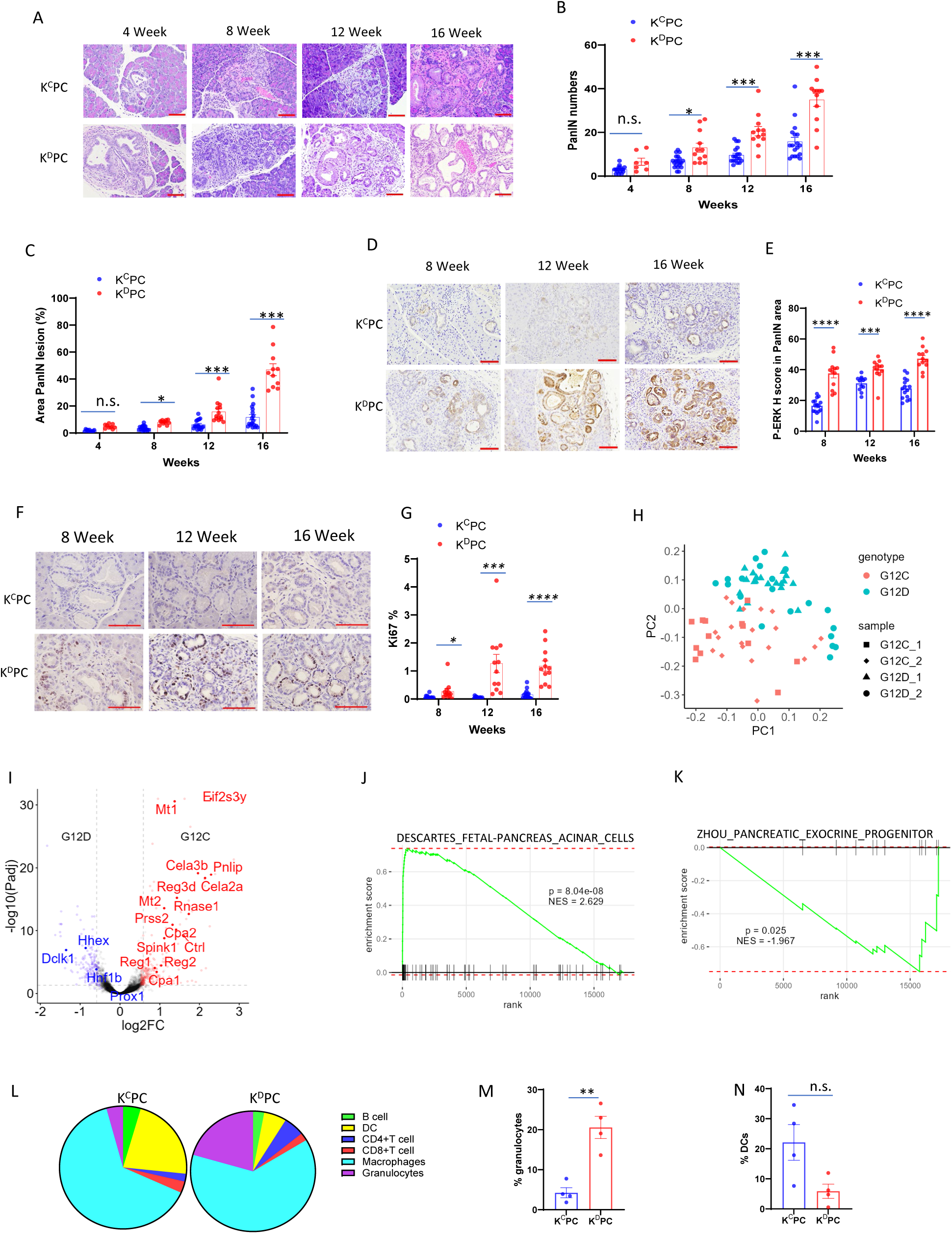
Tumor development is delayed in the K^C^PC model compared to K^D^PC model. **A.** HE staining of mouse pancreas tissues from K^C^PC and K^D^PC model between 4 to 16 weeks. Scale bar: 100 µm. **B.** PanIN/ADM lesions numbers at different weeks of K^C^PC and K^D^PC mice under 200X fields. **C.** PanIN/ADM lesions area percentage at different weeks of K^C^PC and K^D^PC mice under 40X fields. **D. E,** Representative images and H score of p-ERK IHC staining in the PanIN areas of mice tissues at different weeks of K^C^PC and K^D^PC mice. Scale bar: 100 µm. **F. G**, Representative images and percentage of Ki67 IHC staining of mice tissues at different weeks of K^C^PC and K^D^PC. Scale bar: 100 µm. **H.** Principal component analysis (PCA) of top 3000 variedly expressed genes showed a clear difference between K^D^PC and K^C^PC samples. **I.** Volcano plot of DEG analysis showed differentially expressed genes between K^D^PC and K^C^PC samples. The Acinar cell marker genes (red) and progenitor cell marker genes (blue) are labelled. **J.** Gene set enrichment analysis (GSEA) plot showing the enrichment scores for acinar markers in K^C^PC samples. **K.** GSEA plot showing the enrichment scores for exocrine progenitor markers in K^D^PC samples. **L.** Multiplex immunofluorescent staining results showed composition of immune cell populations in K^C^PC and K^D^PC samples. **M.** Quantification of granulocytes populations in total immune cells in K^C^PC and K^D^PC samples. Data, mean ± SEM; *n* = 4. **N.** Quantification of dendritic cell populations in total immune cells in K^C^PC and K^D^PC samples. Data, mean ± SEM; *n* = 4. n.s. not significant. * *P*<0.05, ** *P*<0.01, *** *P*<0.001, \*\*\*\**P*<0.0001

To gain insight into the molecular differences between theses alleles, whole transcriptomic analysis using the GeoMx Digital Spatial Profiler (DSP) platform was conducted on pan-cytokeratin+ (panCK+) epithelial cells from a total of 73 morphologically similar premalignant lesions randomly selected from the pancreatic tissues of 16-week old K^C^PC (33 lesions from 2 mice) and K^D^PC (40 lesions from 2 mice) mice (Supplementary Fig. S1C). As shown in Fig.1H, principal component analysis of the RNA-seq data indicates that, while there is limited heterogeneity between individuals within the same genotype, the molecular features of the preneoplastic lesions from the K^C^PC model can be distinguished from K^D^PC lesions, implicating KRAS allele-specific molecular underpinnings in processes of PDAC initiation. Gene Set Enrichment Analysis (GSEA) indicates enrichment of acinar cell features in G12C lesions accompanied with the significant upregulation of multiple acinar-specific genes compared to G12D lesions (Fig.1I, J; Supplementary Fig.S1D). On the other hand, G12D lesions show significant upregulation of markers of exocrine progenitor cells (Fig.1I, K; Supplementary Fig.S1E), including DCLK1 which has been shown to be a marker for stem-like cells in preinvasive PDAC (36). Moreover, lesions from both models exhibit similar level of ductal cell marker cytokerain-19 (CK19) (Supplementary Fig.S1F). These findings were further confirmed by immunofluorescence staining of pancreatic tissue of 16-week old K^C^PC and K^D^PC models, showing that G12D epithelial lesions completely lose the expression of acinar marker, amylase, which is partially retained in G12C lesions, while DCLK1+ cells are more frequently observed in G12D lesions compared to K^C^PC model (Supplementary Fig.S1G-H). These data indicates that, compared to KRAS^G12D^, KRAS^G12C^ failed to induce complete ADM which is characterized with the transdifferentiating of acinar cells into progenitor-like ductal cells (37).

Tumor microenvironment (TME), in particular, the immune microenvironment plays an important role for KRAS-driven PDAC initiation and progression (38). Multiplex immunofluorescence was conducted to catalog the pancreatic immune infiltration in 16-week-old K^C^PC and K^D^PC mice, when the pancreata contain extensive premalignant lesions, including ADM and PanINs. The dominant immune population in PDAC TME is myeloid cells, including macrophages, neutrophils, monocytes and dendritic cells (39). Indeed, myeloid cells are the majority of CD45^+^ immune cells in the pancreata of both K^C^PC and K^D^PC mice (Fig.1L; Supplementary Fig. S1I). Interestingly, while G12D pancreata contains more CD11b+Ly6g+ granulocytes (Fig. 1L-M; Supplementary Fig. S1I), the pancreata from K^C^PC mice exhibit trend of dendritic cells increasing (Fig.1L, N, *P*=0.069; Supplementary Fig. S1I). Granulocytes are the main components of myeloid derived suppressive cell (MDSC) which plays a major role in the immunosuppressive tumor microenvironments in PDAC (40). This result suggests that the TME of K^C^PC mice is less immunosuppressive than that of K^D^PC model, which may partially explain the slower progression of K^C^PC tumors. Together, our data indicates that the delayed tumor development in the K^C^PC model, compared to K^D^PC model, is correlated with decreased KRAS signaling, incomplete ADM and alterations in the immune microenvironment.

The relatively weak tumor development in K^C^PC model results in significantly prolonged median overall survival compared to K^D^PC model (41 weeks vs 22 weeks) (Fig.2A). Histopathology analysis of advanced tumors from the K^C^PC model revealed similar histological features of ductal adenocarcinoma compared to K^D^PC model or human PDAC (Fig.2B). In addition, PDAC lesions from the K^C^PC and K^D^PC models exhibited similar total KRAS, activated KRAS, phospho-ERK, phospho-AKT and phospho-S6K1 levels (Fig.2C-E; Supplementary Fig.S2A). Bulk RNA-seq was conducted to compare the transcriptomic features between K^C^PC- and K^D^PC-derived tumor cultures. Principle component analysis indicates that the K^D^PC lines are clustered into two groups representing the two major PDAC molecular subtypes (41), with the majority belonging to progenitor subtype and a few lines enriched with squamous subtype signatures (Fig.2F). Interestingly, the K^C^PC lines do not form separate groups, but are clustered together with K^D^PC lines of the progenitor subtype (Fig.2F).

**Figure 2.**
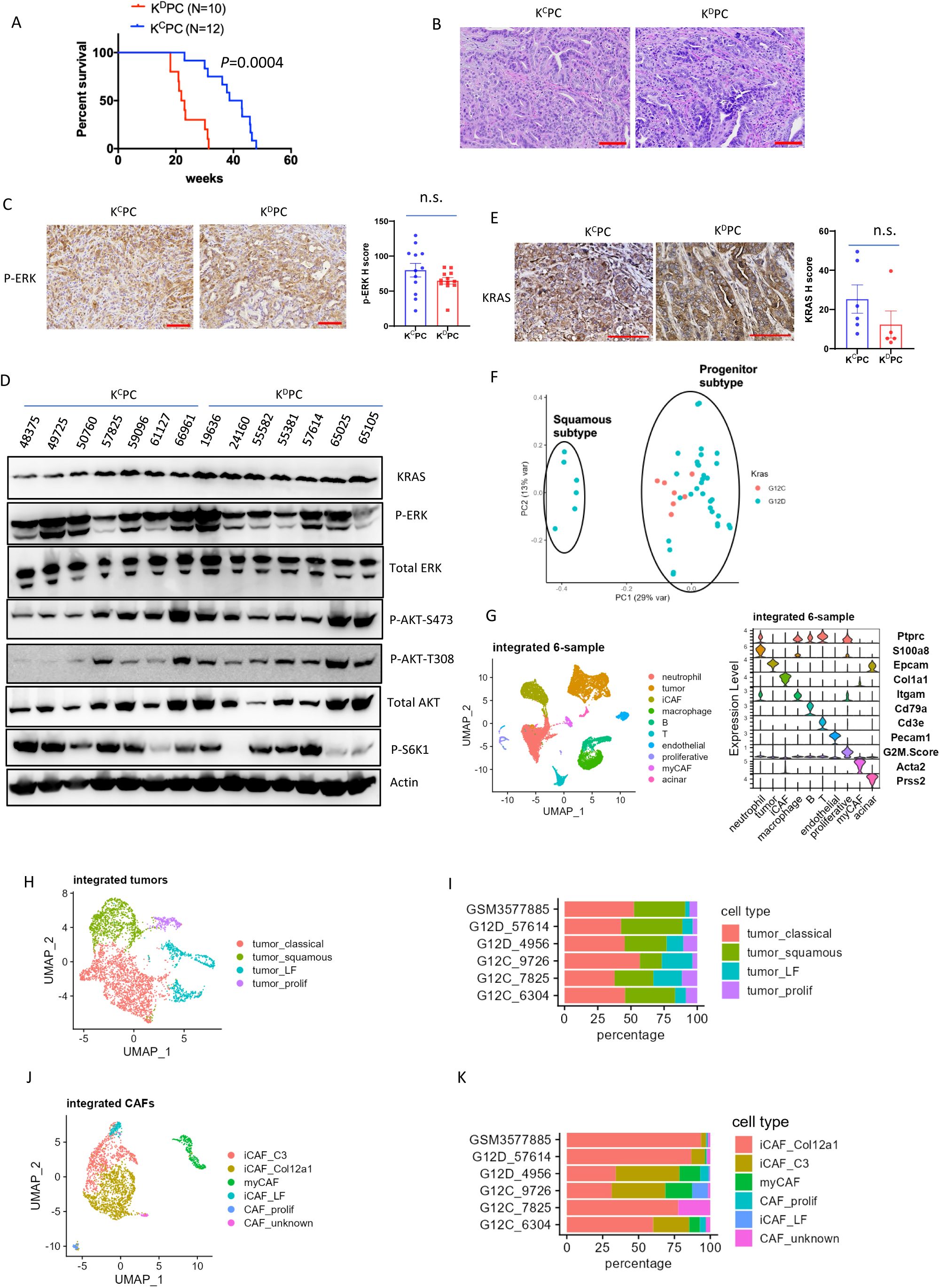
Advanced PDAC from K^C^PC and K^D^PC models exhibit similar histopathological and molecular features. **A**, Kaplan–Meier curves depicting survival of K^C^PC and K^D^PC mice. Statistical significance was determined by the log-rank test. **B**, HE staining of mouse pancreatic tumor tissues from K^C^PC and K^D^PC model. Scale bar: 100 µm. **C**, Representative images and H score of p-ERK IHC staining of mice tumor tissues of K^C^PC and K^D^PC model. Scale bar: 100 µm. n.s. not significant. **D**, Western blots of whole-cell lysates from K^C^PC and K^D^PC cancer cell lines. **E**, Representative images and H score of KRAS IHC staining of mice tumor tissues of K^C^PC and K^D^PC model. Scale bar: 100 µm. n.s. not significant. **F**, PCA of top 5000 variedly expressed genes showed K^D^PC lines are clustered into two groups, and K^C^PC lines are clustered together with K^D^PC lines of the progenitor subtype. **G**, UMAP on three K^C^PC and K^D^PC tumor tissues. Populations are identified by color (see legend), Violin plots of key markers used to define the identified cell populations were shown in right panel. **H**, **I**, Integrated UMAP of cancer cells from K^C^PC and K^D^PC mice showing Louvain clusters (**H**) and tumor subtypes (**I**). **J, K**, Integrated UMAP of cancer-associated fibroblasts (CAFs) from K^C^PC and K^D^PC mice showing Louvain clusters (**J**) and CAF types (**K**).

To more deeply interrogate the cancer cells and associated TME components, single cell RNA sequencing (scRNA-seq) was conducted on advanced tumors from K^C^PC (n=3) and K^D^PC (n=2) models. A recently published scRNA-seq data from the same K^D^PC model was also included for integrated analysis (42). Graph-based population delineation was performed, and populations were assigned an identity based on lineage markers as shown in Fig. 2G. Integrated analysis of all cancer cells from 6 samples identified 4 major clusters (Fig.2H; Supplementary Fig.S2B). While the two major cancer cells clusters are enriched with signatures of the progenitor or squamous subtype, we also identified a small population of highly proliferative cancer cells and a group of squamous subtype cancer cells with low expression feature (LF) (Fig.2H). Notably, while all four cancer cell clusters can be found across the 6 samples, the cancer cell composition exhibit strong inter-tumoral heterogeneity and no significant differences can be identified between the two KRAS genotypes (Fig.2I). As an important component of PDAC TME, cancer-associated fibroblasts (CAFs) comprised 2-30% of total cell populations among the 6 samples (Table S2). In addition to the myofibroblastic CAF (myCAF) cluster with high expression of *Acta2*, *Myl9* and *Tagln* (43), inflammatory CAF (iCAF) is the major component of the CAF population (Fig.2J-K; Supplementary Fig.S2C). Among the iCAF population, we further identified three subclusters, a cluster with high *Col12a1* expression, a cluster with high expression of *complement 3* (*C3*) and *complement 4* (*C4*), and a cluster with low expression feature (Fig.2J-K; Supplementary Fig.S2C). A *C3/C4*-high CAF population was also identified in human PDAC (44), although their function remains unclear. On the other hand, we did not consistently observe the antigen-presenting CAF (apCAF), likely due to the limited cell numbers. Similar to previous report (43), among the CD45+ immune cells, myeloid cells, including macrophages, monocytes, dendritic cells and neutrophils, are the predominant component, ranging from 27-89% with no obvious correlation with KRAS genotypes (Table S3). Although the percentage of T cells among the total cells is higher in K^C^PC tumors compared to K^D^PC tumors (Table S2, *P*=0.03), the proportion of T cells among CD45+ immune cells or the composition of different types of T cells does not show significant difference between the two genotypes (Table S3). No correlation was identified between KRAS genotype and other immune cell types, such as B cells and DC cells, either (Table S3). Together, our data indicates that, while KRAS^G12C^-driven PDAC premalignant lesions exhibit distinctive molecular features compared to the KRAS^G12D^-driven ones, there is no obvious cellular or molecular difference between the two KRAS genotypes once the tumor progresses to an advanced stage.

To evaluate the therapeutic impact of G12Ci in K^C^PC model, allograft tumors were established from two independent primary K^C^PC-derived cultures and treated with AMG 510. While both K^C^PC lines are sensitive to AMG 510 in vitro with IC50 at nanomolar range (Supplementary Fig.S3A), AMG 510 treatment in vivo over the cause of 3 weeks failed to induce obvious tumor regression and instead produced stable or progressive disease based on the RECIST criteria (45) (Fig.3A-B). Recent studies in preclinical models demonstrated that the reprogramming of immune microenvironment, in particular the activation of cytotoxic T cells, is critical for the regression of Kras^G12D^-driven tumors following MRTX1133 treatment (17,46). Therefore, Cytometry by Time of Flight (CyTOF) analysis was conducted to characterize the changes in the immune microenvironment of K^C^PC-derived allograft tumors upon AMG 510 treatment. Similar to the findings in Kras^G12D^-driven PDAC models, the most dominant subset of CD45+ TME cells in the K^C^PC tumors is CD11b^+^Gr-1^+^ myeloid derived suppressive cell (MDSC) population, which decreased significantly following AMG 510 treatment (Fig.3C; Supplementary Fig.S3B). On the other hand, AMG 510 treatment leads to significant increase of CD8^+^ T cells, dendritic cells (DCs) and macrophages (Fig.3C; Supplementary Fig.S3B). Additional analysis of macrophage population further revealed an increase of MHC II^+^ M1-like macrophages and concomitant decrease in ARG1^+^ M2-like macrophages following AMG 510 treatment (Fig.3D; Supplementary Fig.S3C), indicating that KRAS^G12C^ inhibition skews macrophages toward anti-tumor phenotype. CyTOF data was further confirmed by fluorescence-activated cell sorting (FACS) analysis (Supplementary Fig.S3D). IHC of allograft tumors further verified the alterations in immune infiltration, showing increased CD8^+^ T cells and F4/80^+^ macrophages in AMG 510-treated tumors compared to vehicle-treated controls (Supplementary Fig.S3E). Together, these data indicate that AMG 510 treatment engenders an anti-tumor immune profile, reminiscent of the changes in immune populations in KRAS^G12D^-driven tumors treated with MRTX1133 (17,46). However, the lack of obvious KRAS inhibition-induced tumor regression in the K^C^PC model, in contrast to KRAS^G12D^-driven models, implies that cancer cells can sustain viability in a relatively ‘hot’ anti-tumor immune microenvironment.

**Figure 3.**
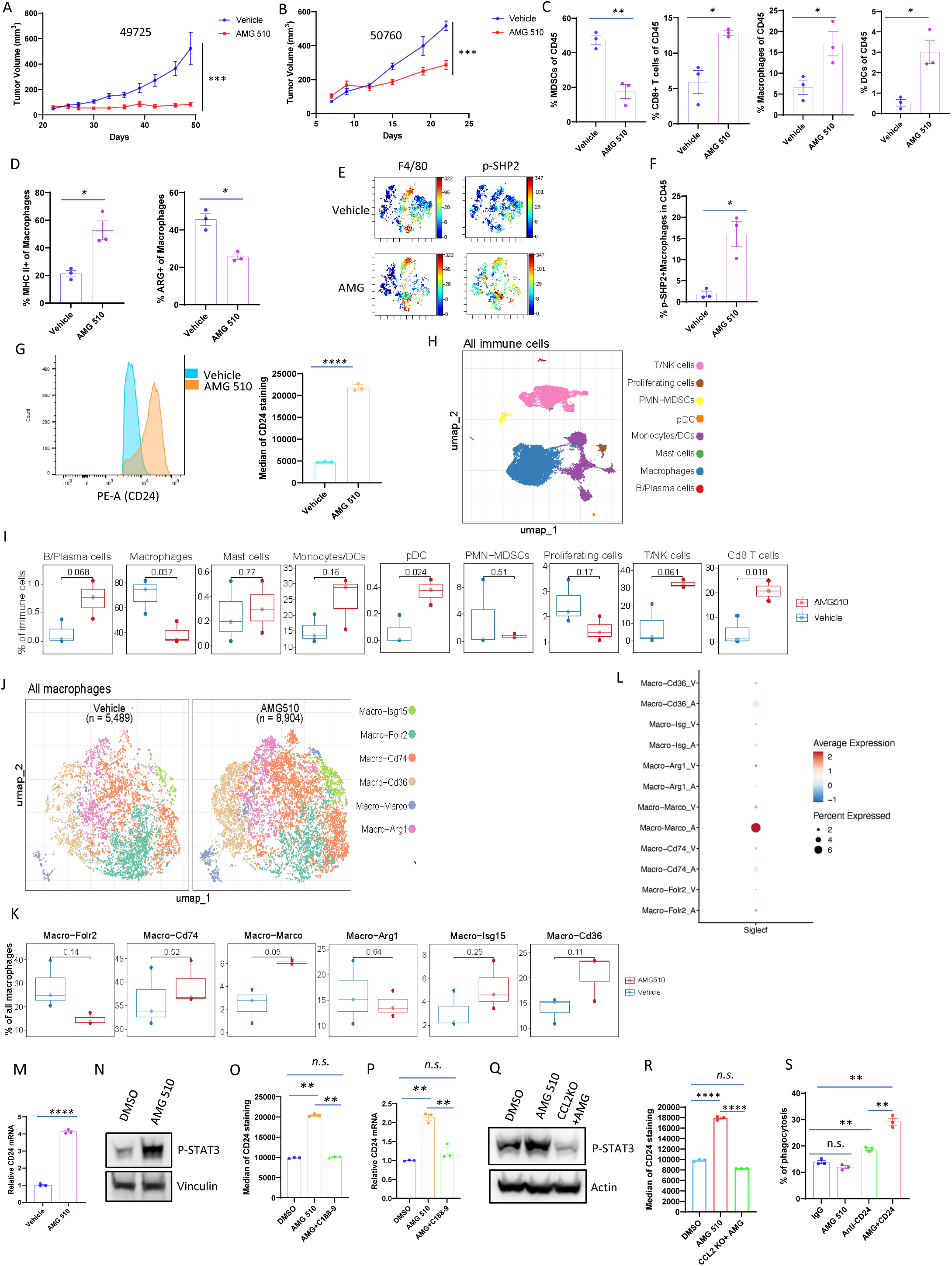
KRAS^G12C^ inhibitor reprograms PDAC tumor microenvironment and upregulates cancer cell CD24 expression. **A, B,** Growth curves of subcutaneous allograft tumors (**A**,49725; **B**, 50760) show change in tumor volume over vehicle or AMG 510 treatment. Data, mean ± SEM. **C,** Quantification of the immune cell populations by CyTOF immune profiling. Data, mean ± SEM; *n* = 3. The statistical difference between vehicle and AMG 510 treated groups was determined by two-tailed *t* tests. **D,** Quantification of macrophage populations by CyTOF analysis. Data, mean ± SEM; *n* = 3. **E**, Representative viSNE plots of p-SHP2+ macrophage populations in K^C^PC mouse pancreatic tumor with vehicle or AMG 510 treatment. **F**, Quantification of p-SHP2+ macrophage populations in **E**. Data, mean ± SEM; *n* = 3. **G**, Median of membrane CD24+ in 50760 K^C^PC cancer cells treated with vehicle or AMG 510 was measured by flow cytometry. Data, mean ± SEM; the statistical difference between experimental groups was determined by two-tailed *t* tests. **H,** UMAP on three vehicle and AMG 510 treated 49725 K^C^PC tumor tissues. Immune cell populations are identified by color (see legend). **I**, Percentages and *P* value of each cell type in **H**. **J,** UMAP on three vehicle and AMG 510 treated 49725 K^C^PC tumor tissues. Subpopulations of macrophages are identified by color (see legend). **K,** Percentages and *P* value of each subtype in **J**. **L,** Expression of Siglecf in different subtypes of macrophages with vehicle or AMG 510 treatment. V: vehicle; A: AMG 510. **M**, qRT-PCR for *CD24* expression in 50760 K^C^PC cancer cells treated with vehicle or AMG 510. Data, mean ± SEM; *n* = 3. The statistical difference was determined by two- tailed *t* tests. **N**, Western blots of whole-cell lysates from vehicle or AMG 510 treated 50760 K^C^PC cancer cell lines. **O**, Median of membrane CD24+ in 50760 K^C^PC cancer cells treated with vehicle, AMG 510 or AMG plus C188-9 was measured by flow cytometry. Data, mean ± SEM; the statistical difference between experimental groups was determined by two-tailed *t* tests. **P**, qRT- PCR for *CD24* expression in 50760 K^C^PC cancer cells treated with vehicle, AMG 510 or AMG plus C188-9. Data, mean ± SEM; *n* = 3. The statistical difference was determined by two- tailed *t* tests. **Q,** Western blots of whole-cell lysates from 50760 K^C^PC cancer cell lines treated with vehicle, AMG 510 or AMG 510 with CCL2 knockout. **R,** Median of membrane CD24+ in 50760 K^C^PC cancer cells treated with vehicle, AMG 510 or AMG 510 with CCL2 knockout was measured by flow cytometry. **S**, In vitro phagocytosis of mouse 50760-GFP PDAC cells cocultured with macrophage cell line RAW264.7 in the presence of AMG 510, anti-CD24 mAb, or dual treatment vs. IgG control. Phagocytosis was measured as the number of F4/80+, GFP+ macrophages, quantified as a percentage of the total F4/80+ macrophages. Data, mean ± SEM; n = 3. Statistics with significance were indicated, n.s. not significant; * P <0.05; ** P <0.01; ***, P < 0.001, **** P <0.0001.

A closer examination of the M1-like macrophages accumulated upon AMG 510 treatment identified significant upregulation of SHP2 phosphorylation (Fig.3E-F), an immune-inhibitory signaling marker(47), suggesting the suppression of macrophage function. Phagocytosis is the major way that macrophages clear cancer cells and cancer cells enabling evasion from phagocytosis via expression of CD47 and CD24 transmembrane proteins which act as anti-phagocytic “don’t eat me” signals (30,48). CD47 or CD24 on cancer cells, by binding to SIRPa or Siglec-10 (Siglec-E, -F, -G in mouse), respectively, recruits SHP1 or SHP2, and inhibits phagocytosis of macrophages (30,48). In accordance with the increase in phospho-SHP2 level in macrophages, CyTOF analysis demonstrates that CD24 intensity is upregulated in cancer cells after AMG 510 treatment (Supplementary Fig.S3F). FACS analysis in two independent K^C^PC tumor lines further confirmed that CD24, but not CD47, is upregulated upon KRAS^G12C^ inhibition (Fig. 3G; Supplementary Fig.S3G-I).

To further characterize the immune reprogramming in response to the KRAS^G12C^ inhibitor, scRNA-seq was conducted on CD45+ enriched immune cells from vehicle (n=3) and AMG 510 (n=3) treated syngeneic orthotopic allograft K^C^PC tumors. Integrated analysis of all immune cells from 6 samples identified 8 major clusters (Fig. 3H). Myeloid cells, including macrophages, monocytes, dendritic cells and MDSC, are the predominant components (Fig. 3H). Consistent with the CyTOF data, AMG 510 treatment leads to significant increase of CD8^+^ T cells and p-DCs (Fig. 3I). Macrophages were further assigned to six clusters, including Folr2+, Cd74+, Marco+, Arg1+, Isg15+ and Cd36+ macrophages (Fig. 3J). We found that Marco+ macrophage was significantly increased after AMG 510 treatment (P=0.05) (Fig. 3K). There was no significant difference in other clusters of macrophages (Fig. 3K). Interestingly, Siglecf, one of the CD24 receptors, was highly expressed in Marco+ macrophages and was further induced after AMG 510 treatment (Fig. 3L).

QRT-PCR analysis indicates that the upregulation of CD24 after AMG 510 treatment is at transcription level (Fig.3M). To further interrogate the mechanisms underlying CD24 upregulation, kinase array analysis is conducted following AMG 510 treatment in K^C^PC cells as the immediate downstream signaling of KRAS oncogene is largely mediated by phosphorylation events. As expected, AMG 510 treatment results in the inhibition of multiple phosphorylation events, including the decrease of phosphorylation of MAPK and its downstream target c-Jun (Supplementary Fig.S3J). Intriguingly, STAT3 phosphorylation at tyrosine^705^ (Y^705^), a marker for STAT3 activation (49), is increased in AMG 510-treated cells (Fig.3N; Supplementary Fig.S3J). Pre-treatment with STAT3 inhibitor C188-9 prevented the AMG 510-induced upregulation of CD24 (Fig. 3O-P; Supplementary Fig.S3K), indicating that STAT3 activation is required for the induction of CD24 in cancer cells upon KRAS^G12C^ inhibition. STAT3^Y705^ becomes phosphorylated in response to cytokines and growth factors. Interestingly, cytokine array analysis discovered that CCL2 was significantly increased in K^C^PC cells following AMG 510 treatment (Supplementary Fig.S3L). CCL2 is also known as monocyte chemoattractant protein-1 (MCP-1), which is a potent chemokine for monocytes. CCL2 has been shown to activate STAT3 through its receptor CCR2 (50). CCL2 knockout or treatment with CCL2 neutralizing antibody prevented AMG510-induced STAT3 phosphorylation (Fig.3Q; Supplementary Fig.S3M) and CD24 upregulation (Fig. 3R; Supplementary Fig.S3N), indicating that CCL2 is required for the activation of STAT3 and induction of CD24 in cancer cells upon KRAS^G12C^ inhibition. K^C^PC cell and macrophage co-culture was established to further investigate whether the induction of CD24 in cancer cells indeed suppresses macrophage phagocytic function. As shown in Fig.3S, while AMG 510 treatment alone fails to affect macrophage phagocytosis, treatment of anti-CD24 monoclonal antibody (mAb) to block CD24-Siglec-G interaction increases macrophage phagocytosis which is further significantly enhanced by co-treatment with AMG 510.

To further evaluate translation relevance of targeting CD24, sub-cutaneous syngeneic allograft models were established with K^C^PC lines for treatment with anti-CD24 mAb alone or in combination with AMG 510. While treatment with either AMG 510 or anti-CD24 mAb alone inhibits tumor growth, the combination further slows down the tumor growth and leads to tumor regression in some cases (Fig. 4A-B). The cooperation between co-targeting KRAS^G12C^ and CD24 was further confirmed in syngeneic orthotopic allograft models. Compared with AMG 510 or anti- CD24 mAb alone, combination therapy further decreased tumor volume (Fig.4C) and significantly prolonged mice overall survival (Fig. 4D). The benefit of combination therapy was also confirmed in GEM model (Fig. 4E; Supplementary Fig. S4A). Moreover, while anti-CD24 mAb treatment alone failed to directly inhibit cancer cell proliferation (Supplementary Fig. S4B), the anti-tumor effect of AMG 510 and anti-CD24 mAb combination is accompanied with significant increase in macrophage phagocytosis (Fig.4F) and decrease of phospho-SHP2 (Fig.4G) in the allograft tumors, suggesting the activation of macrophage function upon combination treatment. To further determine the requirement of macrophages for the anti-tumor effect of the combination therapy, tumor macrophages are depleted with anti-CSF1R mAb treatment in the K^C^PC allograft models (Supplementary Fig.S4C). As shown in Fig.4H, depletion of macrophages completely reverses the tumor growth inhibition induced by the treatment of AMG 510 and anti-CD24 mAb. On the other hand, the anti-tumor effect of the combination therapy is not affected in TCR-/- mice (Fig. 4I). Together, our data indicates the essential role of macrophage-mediated innate immunity, rather than T-cell-mediated adaptive immunity, for the therapeutic effect of AMG 510 in combination with anti-CD24 mAb.

**Figure 4.**
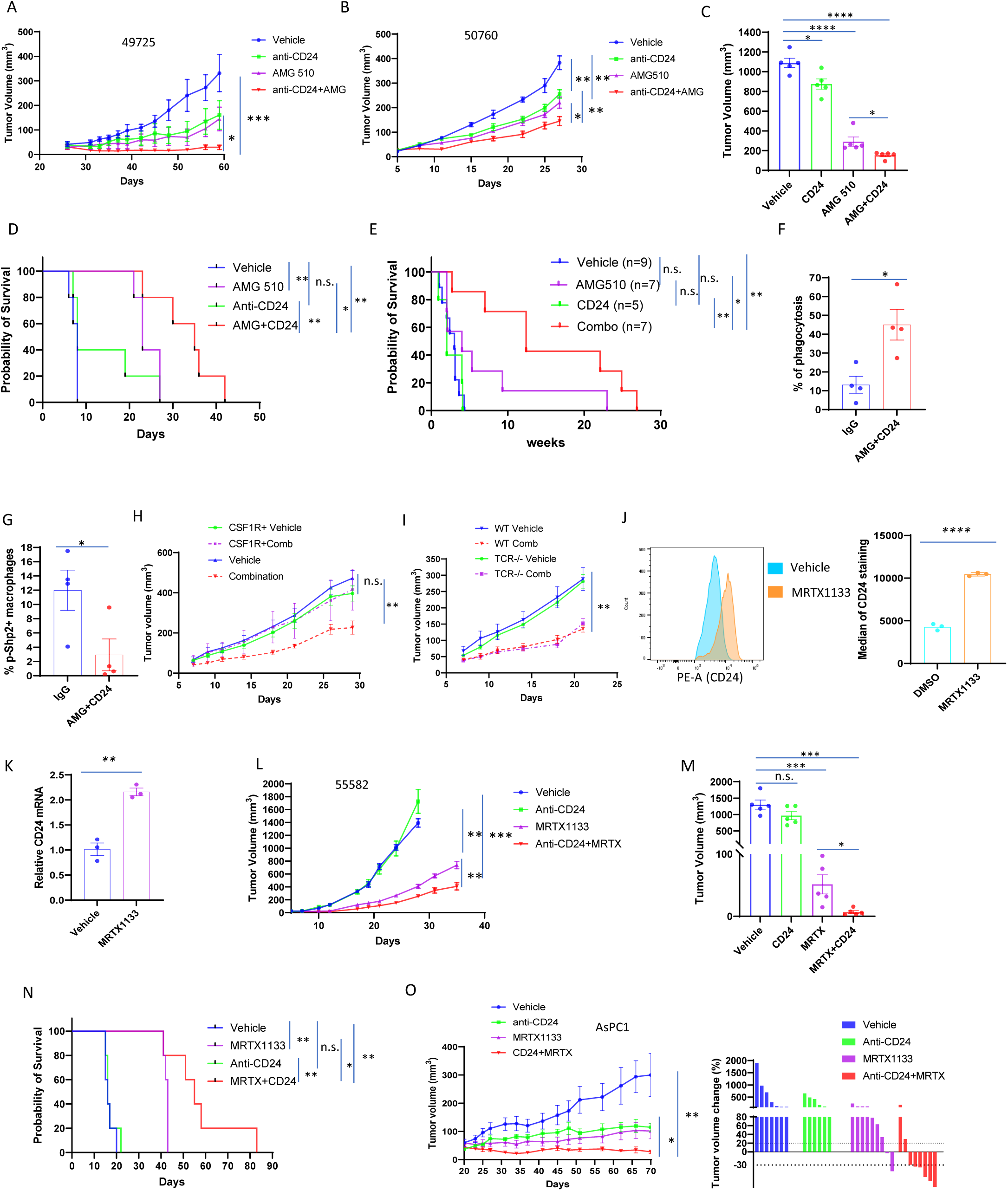
Targeting CD24 sensitizes pancreatic cancer to both KRAS^G12C^ *and* KRAS*^G12D^* inhibitors in vivo. **A, B,** Growth curves of subcutaneous allograft K^C^PC tumors (**A**,49725; **B**, 50760) show change in tumor volume over vehicle, anti-CD24, AMG 510 or anti-CD24 + AMG 510 treatment. Data, mean ± SEM. **C,** Orthotopically injected K^C^PC tumors (50760) were measured by MRI for tumor volumes in vehicle, anti-CD24, AMG 510 or anti-CD24 + AMG 510 treatment. Data, mean ± SEM. **D,** Kaplan–Meier curves depicting overall survival of K^C^PC mice in **C**. Statistical significance was determined by the log-rank test. **E**, Kaplan–Meier curves depicting overall survival of K^C^PC GEM mice with vehicle, anti-CD24, AMG 510 or anti-CD24 + AMG 510 treatment. Statistical significance was determined by the log-rank test. **F**, In vivo phagocytosis. Mouse 50760-GFP PDAC cells were injected into the right flank of C57BL/6 mice and treated with IgG control or AMG 510+ anti-CD24 mAb for 14 days. Mice tumor tissues were collected for flow cytometry. Phagocytosis was measured as the number of F4/80+, GFP+ macrophages, quantified as a percentage of the total F4/80+ macrophages. Data, mean ± SEM; *n* = 4. **G**, Mice tumor tissues collected in **F** were measured by flow cytometry for F4/80 and p-SHP2. The percentage of p-SHP2+ macrophages were compared in IgG control or AMG 510+ anti-CD24 mAb groups. Data, mean ± SEM; n = 4. **H**, Growth curves of subcutaneous allograft K^C^PC tumor (50760) show changes in tumor volume over vehicle, vehicle + anti-CSF1R, anti-CD24 + AMG 510 (combination), or anti-CSF1R + combination treatment. Data, mean ± SEM. **I,** Growth curves of subcutaneous allograft K^C^PC tumor (50760) in C57BL/6 mice or TCR-/- mice show changes in tumor volume over vehicle and anti-CD24 + AMG 510. Data, mean ± SEM. **J**, Median of membrane CD24+ in K^D^PC cancer cells (55582) treated with vehicle or MRTX1133 was measured by flow cytometry. Data, mean ± SEM; the statistical difference between experimental groups was determined by two-tailed *t* tests. **K**, qRT-PCR for *CD24* expression in K^D^PC cancer cells (55582) treated with vehicle or MRTX1133. Data, mean ± SEM; *n* = 3. The statistical difference was determined by two-tailed *t* tests. **L**, Growth curves of subcutaneous allograft K^D^PC tumor (55582) show change in tumor volume over vehicle, anti-CD24, MRTX1133 or anti-CD24 + MRTX1133 treatment. Data, mean ± SEM. **M**, Orthotopically injected K^D^PC tumors (55582) were measured by MRI for tumor volumes in vehicle, anti-CD24, MRTX1133 or anti-CD24 + MRTX1133 treatment. Data, mean ± SEM. **N,** Kaplan–Meier curves depicting overall survival of K^D^PC mice in **M**. Statistical significance was determined by the log-rank test. **O**, **Left**, Growth curves of human KRAS G12D tumors (AsPC1) show change in tumor volume over vehicle, anti-CD24, MRTX1133 or anti-CD24 + MRTX1133 treatment. Data, mean ± SEM. **Right,** Waterfall plot of vehicle, anti-CD24, MRTX1133 or anti-CD24 + MRTX1133-treated tumors showing a change in tumor volume after treatment compared with baseline at day 0. Each bar represents a single tumor. Statistics with significance were indicated, n.s. not significant; * P <0.05; ** P <0.01; ***, P < 0.001, **** P <0.0001.

The most common KRAS mutation in human PDAC is KRAS^G12D^ which is present in approximately one-third of PDAC patients (5). We next investigated whether CD24-mediated macrophage inhibition also occurs in KRAS^G12D^-driven PDAC following KRAS inhibition. Similar to the observation from K^C^PC cells, treatment with KRAS^G12D^ inhibitor, MRTX1133 (16), also significantly induces the transcription and cell surface level of CD24 in mouse K^D^PC cancer cells (Fig.4J-K). Importantly, KRAS^G12D^ inhibition with MRTX1133 treatment also upregulates CD24 expression in multiple patient-derived xenografts (PDX) model (Supplementary Fig.S4D). Similar to the mechanisms identified in the K^C^PC model, MRTX1133 treatment also upregulated CCL2 expression and induced STAT3 phosphorylation (Supplementary Fig.S4E-F) in K^D^PC cells. Blocking CCL2 with anti-CCL2 reversed MRTX1133 induced CD24 expression (Supplementary Fig.S4G-H). Importantly, treatment with anti-CD24 mAb further enhances the anti-tumor effect of MRTX1133 in both subcutaneous (Fig. 4L) and orthotopic (Fig. 4M; Supplementary Fig.S4I) allograft model of K^D^PC tumors. Moreover, combination therapy in orthotopic allograft model significantly prolongs mice overall survival (Fig. 4N). Furthermore, anti-CD24 mAb in combination with MRTX1133 induces tumor regression in human pancreatic cancer cell line AsPC1-derived xenograft tumors in nude mice (Fig.4O). Together, our data indicates that targeting CD24 enhances macrophage phagocytosis and sensitizes KRAS-driven PDAC to KRAS targeted therapies.

## Discussion

While the genetics of KRAS mutations in PDAC have been well-established(4,5), how these different oncogenic alleles exert common or distinct functions in PDAC biology is not well understood. Recent studies in preclinical models revealed that KRAS^G12R^, a rare KRAS mutation uniquely enriched in PDAC, is defective in PI3K signaling and exhibit weaker oncogenesis in PDAC compared to G12D mutation (31,51), shedding light on the allele-specific biology of KRAS mutations. Here, our data indicates that, compared to KRAS^G12D^, KRAS^G12C^ leads to relatively weaker activation of the downstream MAPK signaling in premalignant lesions, which results in the delay in tumor progression in the p53 deficient GEM model. The delayed PDAC initiation observed in our K^C^PC model is consistent with previous study from the p48Cre, Kras^G12C^ ^L/+^ model (KC-G12C) (52). However, it has been shown that CRISPR-mediated p53 deletion in pancreatic ductal organoid derived from the KC-G12D (p48Cre, Kras^G12D^ ^L/+^) or KC-G12C model results in similar allograft tumor development in immunodeficient nude mice (31). In contrast, by using the autochthonous K^C^PC model with heterozygous p53 deletion, our data demonstrated that malignant tumor progression is also delayed in Kras^G12C^-driven PDAC compared to those tumors with Kras^G12D^. Similar observations were also recently reported in KRAS^G12C^-driven lung cancer model (53).

The decreased MAPK signaling downstream of KRAS^G12C^ could be correlated with the unique biochemistry feature of the G12C mutation, which has been shown to maintain largely intact intrinsic GTPase activity (3). Recent study also revealed that KRAS^G12C^ exhibit distinctive phospholipid binding profile compared to KRAS^G12D^ (54), which may affect its localization in membrane nanodomains and thus impact the downstream signaling. In addition, inflammation has been shown to play a critical role in activation of KRAS signaling during PDAC initiation (55). Our data demonstrates that the immune environment in KRAS^G12C^-induced premalignant lesions shows differences from that of KRAS^G12D^ lesions, which may also contribute to the differential KRAS signaling observed in these two models. Notably, although the tumor development is delayed in the K^C^PC model compared to K^D^PC model, the advanced tumors from these models exhibit same pathology features, comparable level of MAPK signaling and similar molecular features. This implicates the involvement of additional genetic or epigenetic events for the full activation of KRAS^G12C^ during tumor progression. Interestingly, recent analysis of colon and lung cancer samples revealed distinctive co-mutation patterns among different KRAS oncogenes, including the G12C mutation (56), pointing to the need for additional genomic analysis of co-mutations in PDAC with KRAS^G12C^.

Emerging evidence suggests that the immune system may play a critical role in defining the response to KRAS-targeted therapy. Targeting KRAS oncogene in preclinical models of NSCLC and PDAC has been shown to reverse the suppressive immune microenvironment and activate anti-tumor immunity, which plays a critical role for the tumor regression (17,57). However, while the reprogramming of immune microenvironment is critical for the augmentation of anti-tumor effect elicited by KRAS targeted therapy, it may also negatively impact the durability of therapy response through the accumulation of certain immune populations, especially the macrophages. Our data indicates that myeloid cells remain as the predominant immune cell population in the TME upon KRAS targeted therapy, implicating they may play major role in regulating tumor immunity. Indeed, it was recently shown that the recruitment of macrophage into the tumor microenvironment following genetic inactivation of Kras^G12D^ in advanced mouse PDAC promote the survival of cancer cells and bypass of KRAS-dependence (27). Here, our data demonstrated that the anti-tumor function of macrophages in the TME is suppressed by the induction of CD24 in cancer cells following KRAS targeted therapy, undermining the therapeutic benefit of KRAS inhibitors. While more studies are needed to further dissect the macrophage-mediated regulation of innate and adaptive immunities following KRAS inhibition, such as the impact on NK and CD8^+^ T cells, our study provides additional rationale for the co-targeting of immune microenvironment to achieve sustainable therapeutic response to KRAS inhibition in PDAC.

## Acknowledgments

This study was funded by NIH R01CA214793 and The MD Anderson Cancer Center SPORE in Gastrointestinal Cancer DRP Grant P50 CA221707 (to H. Ying), NIH P01CA117969 (to H. Ying, H. Wang, and R.A. DePinho), NIH 1R01CA272744 and ACS RSG-22-017-01-CCB (to W. Yao), The MD Anderson Cancer Center SPORE in Gastrointestinal Cancer CEP Grant P50 CA221707 (to Y. Wei), Andrew Sabin Family Fellows Award (to H. Ying), CPRIT Training Award RP210028 (to E. Y. Yen) and The MD Anderson Cancer Center Support Grant NIH P30 CA016672. This work was funded in part by Odyssey fellowship and Ya-Yen Lee’s fellowship from The University of Texas MD Anderson Cancer Center (to C. F. Li). We acknowledge The MD Anderson Cancer Center Flow Cytometry and Cellular Imaging Core Facility, Research Histology Core Laboratory and Advanced Technology Genomics Core (ATGC). Next generation sequencing was done at Texas Scientific LLC (Houston) with AVITI System from Element Biosciences. We thank Dr. Anirban Maitra for the insightful suggestions during the execution of the experiments and writing of the manuscript. We are thankful to members of the Ying and Yao laboratories for their comments and suggestions.

## Authors’ Contributions

**Y. Wei:** Conceptualization, formal analysis, validation, investigation, visualization, methodology, writing–original draft, writing–review and editing. **M. Liu:** Formal analysis, validation and investigation. **E. Y. Yen:** Formal analysis, methodology, validation and investigation. **J. Yao:** Formal analysis, methodology, validation and investigation. **Z. Xun**: Formal analysis, validation and investigation. **P.T. Nguyen:** Formal analysis, validation and investigation. **X. Wang:** Methodology, validation and investigation. **Z. Yang:** Validation and investigation. **A. Yousef:** Validation and investigation. **D. Pan:** Methodology, validation and investigation. **Y. Jin:** Methodology, validation and investigation. **C.F. Li:** Validation and investigation. **M.S. Theardy:** Validation and investigation. **J. Park:** Validation and investigation. **Y. Cai:** Validation and investigation. **M. Takeda:** Validation and investigation. **M. Vasquez:** Formal analysis, methodology and investigation. **E. M. Park:** Methodology and investigation. **D. H. Peng:** Resources and investigation. **Y. Zhou:** Supervision and investigation. **H. Zhao:** Supervision and investigation. **T.P. Heffernan:** Resources and supervision. **A. Viale:** Supervision and investigation. **H. Wang:** Supervision, resources and investigation. **S.S. Watowich**: Supervision and investigation. **H. Liang:** Supervision and investigation. **D. Zhao:** Resources and investigation. **R.A. DePinho:** Supervision, resources, visualization, writing–review and editing. **W. Yao:** Conceptualization, resources, formal analysis, supervision, funding acquisition, validation, investigation, visualization, methodology, writing–original draft, project administration, writing– review and editing. **H. Ying:** Conceptualization, resources, formal analysis, supervision, funding acquisition, validation, investigation, visualization, methodology, writing–original draft, project administration, writing–review and editing.

## Authors’ Disclosures

R.A. DePinho is Founder and Advisor of Tvardi Therapeutics, Asylia Therapeutics, Stellanova Therapeutics, Nirogy Therapeutics, and Sporos Bioventures. T.P. Heffernan receives advisory fees from Cullgen Inc., Psivant Therapeutics, Isomorphic Labs, Quotient Therapeutics, and Proxygen. No disclosures were reported by the other authors.

---

**Supplementary Figure S1.**
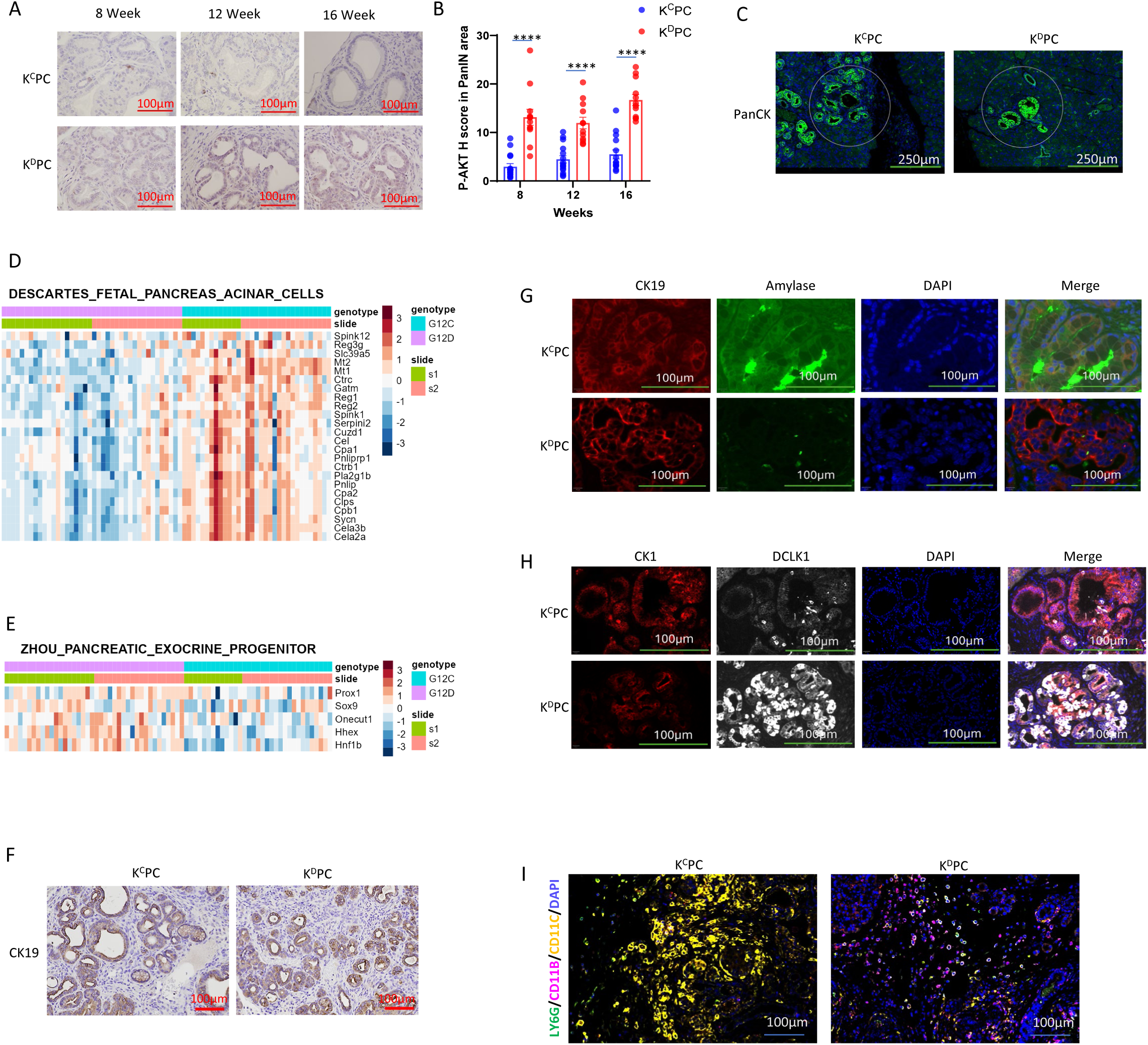
**A. B,** Representative images and H score of p-AKT-S473 IHC staining in the PanIN areas of mice tissues at different weeks of K^C^PC and K^D^PC mice. Scale bar: 100 µm. **** P <0.0001. **C,** AOIs selections for GeoMx digital spatial profiler RNA profiling analysis of K^C^PC and K^D^PC PanIN lesions. For each section, 13-20 AOIs of 200 µm in diameter were selected based on PanCK+ expression. Scale bar: 250 µm. **D,** GSEA term heatmap showed acinar markers are up in K^C^PC samples. **E,** GSEA term heatmap showed exocrine progenitor markers are up in K^D^PC samples. **F,** Representative images of CK19 IHC staining of mice pancreas tissues at 16 weeks of age of K^C^PC and K^D^PC. Scale bar: 100 µm. **G**, Immunofluorescent staining of CK19 and amylase in mice pancreas tissues at 16 weeks of age of K^C^PC and K^D^PC. Nuclei were counterstained with DAPI. Scale bar: 100 µm. **H**, Immunofluorescent staining of CK19 and DCLK1 in mice pancreas tissues at 16 weeks of age of K^C^PC and K^D^PC. Nuclei were counterstained with DAPI. Scale bar: 100 µm. **I.** Representative multiplex immunofluorescent staining images in mice pancreas tissues at 16 weeks of age of K^C^PC and K^D^PC. Scale bar: 100 µm.

**Supplementary Figure S2.**
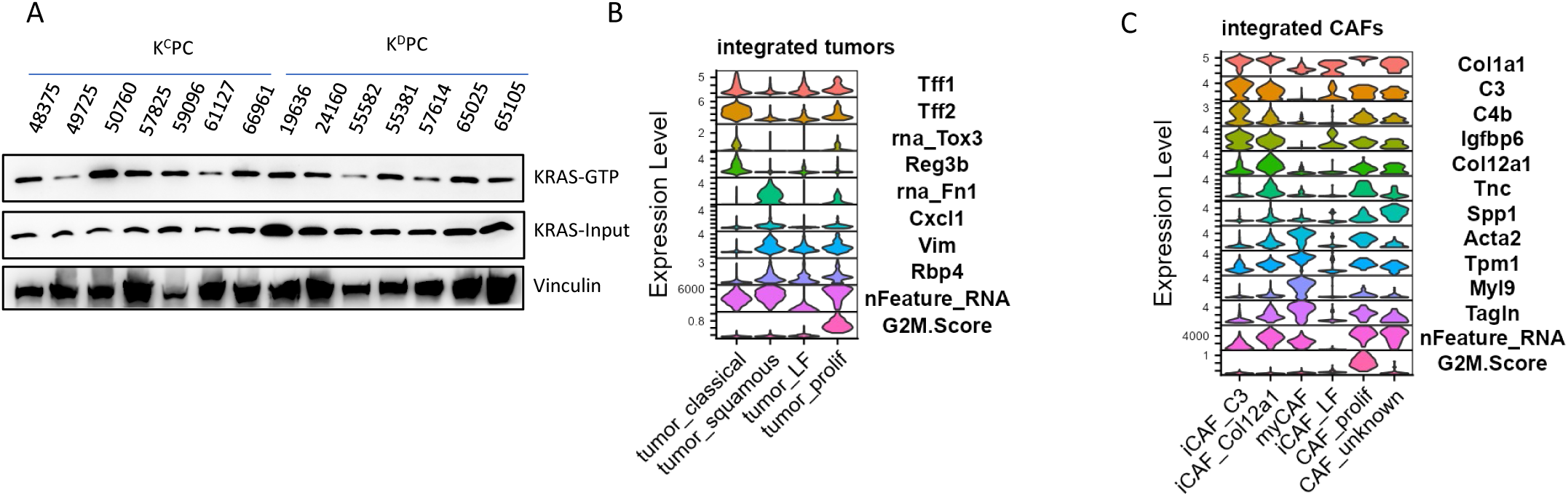
**A**, Ras-GTP levels in K^C^PC and K^D^PC cancer cell lines were assessed by Active Ras Pull-Down and detected by Western blot analysis using mouse monoclonal anti-KRAS antibody. **B**, Violin plots of key markers used to define the identified cancer cell populations. **C**, Violin plots of key markers used to define the identified CAF cell populations.

**Supplementary Figure S3.**
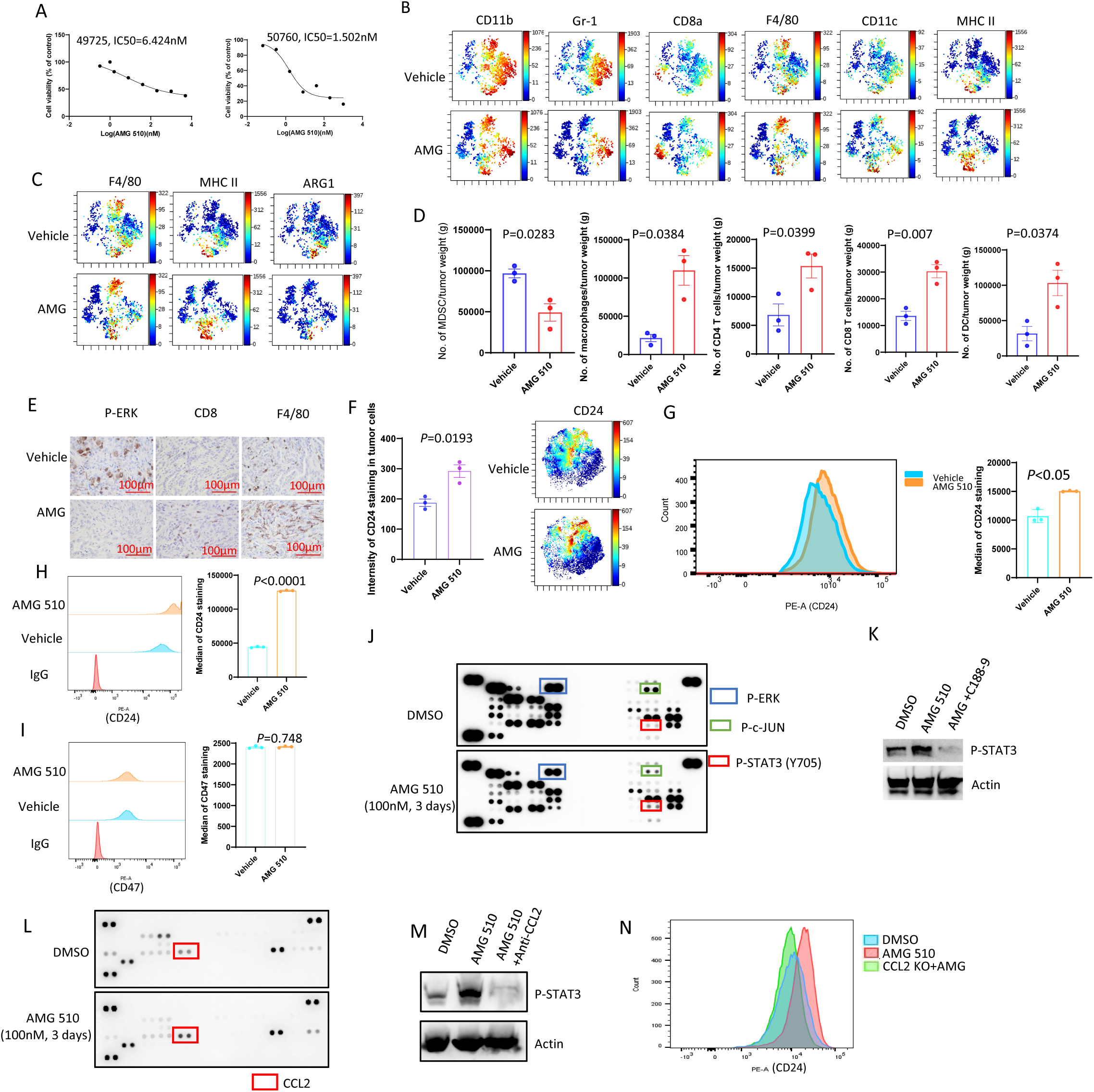
**A,** IC_50_ values for AMG 510 in two mice K^C^PC cell lines was determined by MTT assay. **B**, CyTOF immune profiling by viSNE of K^C^PC mouse pancreatic tumors with vehicle or AMG 510 treatment. **C**, Representative viSNE plots of macrophage populations in K^C^PC mouse pancreatic tumor with vehicle or AMG 510 treatment. **D,** Quantification of the immune cell numbers per tumor tissue gram by FACS in K^C^PC mouse pancreatic tumor with vehicle or AMG 510 treatment. **E,** IHC of allograft tumors showing phospho-ERK was reduced whereas CD8^+^ T cells and F4/80^+^ macrophages were increased in AMG 510-treated tumors compared to vehicle-treated group. **F,** CyTOF analysis demonstrates that CD24 intensity was increased in cancer cells after AMG 510 treatment. **Left**, quantification of CD24 intensity. **Right**, representative viSNE plots of CD24 staining in tumors with vehicle or AMG 510 treatment. **G,** Median of membrane CD24+ in 49725 K^C^PC cancer cells treated with vehicle or AMG 510 was measured by flow cytometry. **H, I,** Median of membrane CD24+ (**H**) or CD47+ (**I**) in 50760 K^C^PC cancer cells treated with vehicle or AMG 510 was measured by flow cytometry. **J,** Kinase array analysis indicated that AMG 510 treatment in 50760 K^C^PC cancer cells results in decrease of p-ERK and p-c-Jun and increase of p-STAT3 (Y705). **K,** Western blots of whole-cell lysates from vehicle, AMG 510 or AMG 510 plus C188-9 treated 50760 K^C^PC cancer cell lines. **L,** Cytokine array analysis indicated that AMG 510 treatment in 50760 K^C^PC cancer cells leads to increasement of CCL2. **M,** Western blots of whole-cell lysates from vehicle, AMG 510 or AMG 510 plus anti-CCL2 antibody treated 50760 K^C^PC cancer cell lines. **N,** Membrane CD24 staining in 50760 K^C^PC cancer cells treated with vehicle, AMG 510, or AMG 510 with CCL2 knockout was measured by flow cytometry.

**Supplementary Figure S4.**
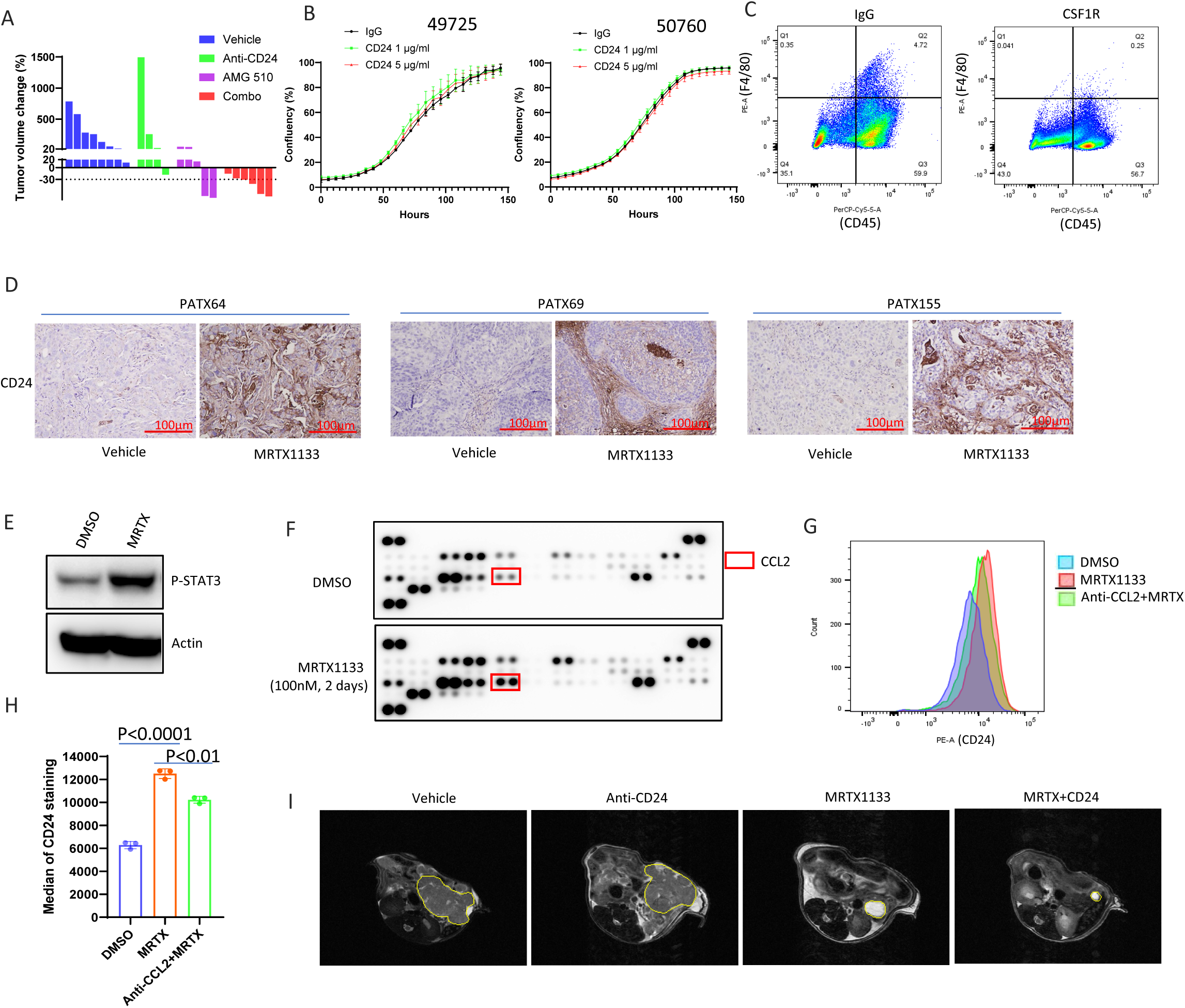
**A,** Waterfall plot of K^C^PC GEM mice tumor with vehicle, anti-CD24, AMG 510 or anti-CD24 + AMG 510 treatment showing a change in tumor volume after treatment compared with baseline at day 0. Each bar represents a single tumor. **B,** Growth curve of two mice K^C^PC cell lines treated with IgG control or anti-CD24 antibody was measured by IncuCyte automatic cell count software. **C,** Macrophage depletion by anti-CSF1R antibody. Mice received 400 μg of anti-CSF1R antibody i.p., every other day whereas control mice were given isotype IgG control. Tumor tissues were collected to confirm macrophage depletion via flow cytometry. Data showed both PE-F4/80+ and PerCP5.5-CD45+ macrophages were 0.25% in CSF1R treated group, compared to 4.72% in IgG treated group. **D**, IHC staining of PDX tumors showing CD24 increased in MRTX1133-treated tumors compared to vehicle-treated tumors. **E,** Western blots of whole-cell lysates from vehicle or MRTX1133 treated 55582 K^D^PC cancer cell lines. **F,** Cytokine array analysis indicated that MRTX1133 treatment in 55582 K^D^PC cancer cells results in an increase of CCL2. **G, H,** Median of membrane CD24+ in K^D^PC cancer cells (55582) treated with vehicle, MRTX1133 or MRTX1133 plus anti-CCL2 antibody was measured by flow cytometry. Data, mean ± SEM. **I,** Representative MRI images of orthotopically injected K^D^PC tumors (55582) with vehicle, anti-CD24, MRTX1133 or anti-CD24 + MRTX1133 treatment. Tumor shape was outlined.

**Supplementary Table S1.**
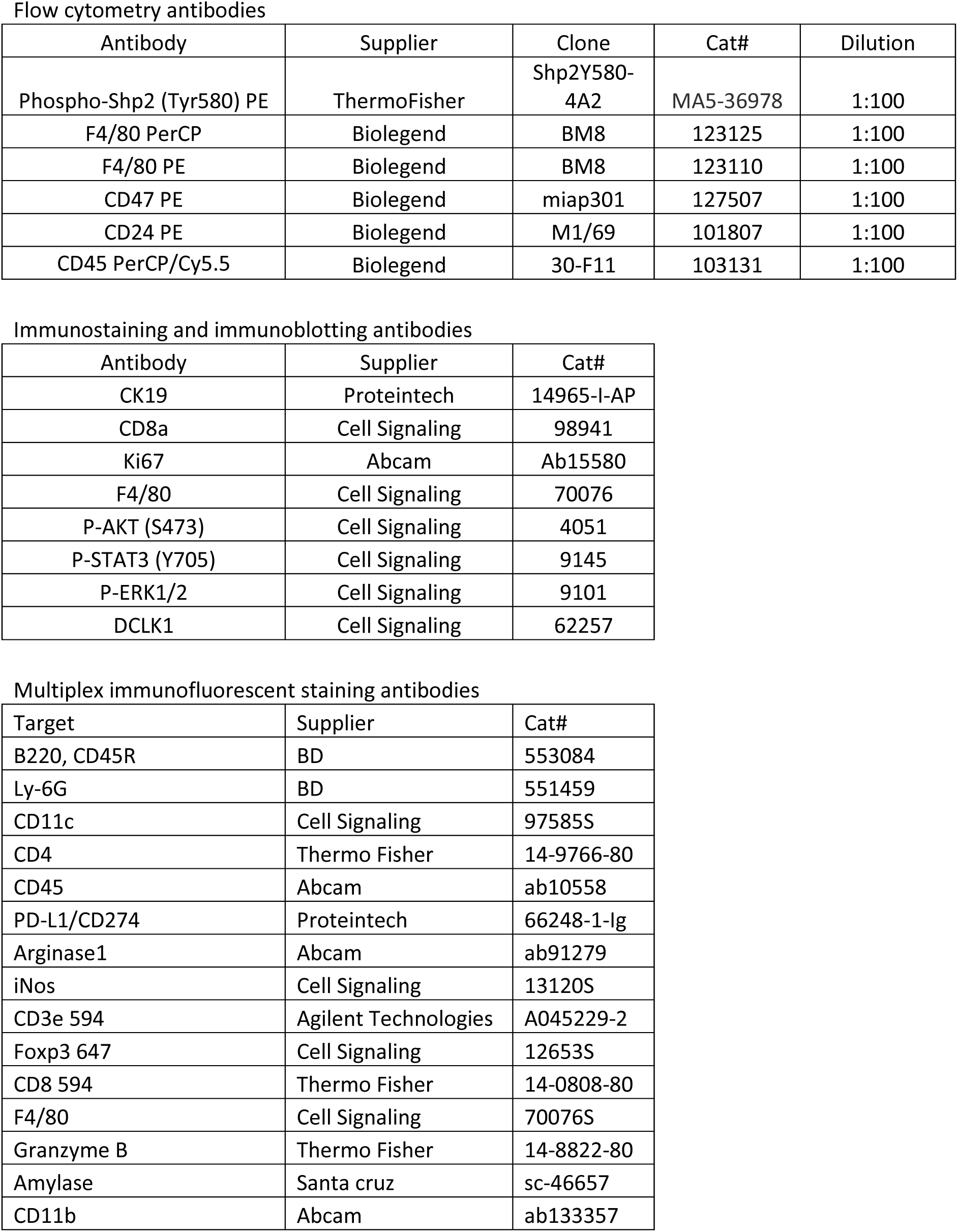
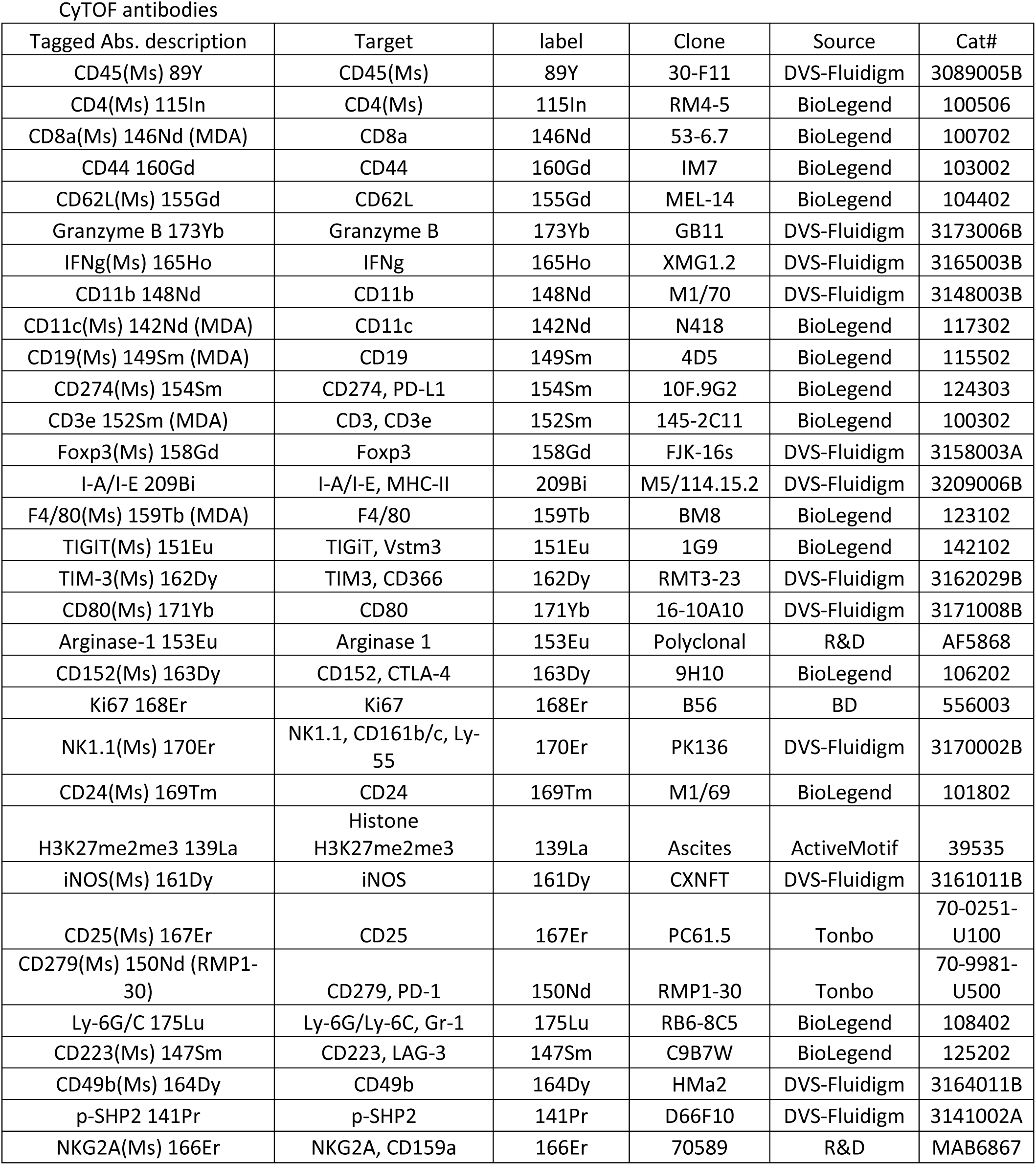
Antibodies list.

**Supplementary Table S2.** Percentage of different cell types in mouse KRAS G12C and G12D tumor tissues.

| Annotation | G12C_6304 | G12C_7825 | G12C_9726 | G12D_4956 | G12D_57614 | GSM3577885 |
| --- | --- | --- | --- | --- | --- | --- |
| Neutrophil | 40.46% | 70.59% | 9.92% | 9.71% | 20.00% | 13.29% |
| Tumor | 17.16% | 5.99% | 40.82% | 33.48% | 43.06% | 16.85% |
| iCAF | 14.59% | 0.96% | 16.62% | 29.83% | 21.73% | 6.64% |
| Macrophage | 8.98% | 8.43% | 1.06% | 5.82% | 4.28% | 45.63% |
| B | 3.60% | 4.89% | 9.48% | 5.71% | 5.61% | 10.84% |
| T | 3.80% | 7.23% | 9.60% | 0.80% | 2.45% | 1.64% |
| Endothelial | 8.37% | 0.33% | 3.06% | 8.11% | 0.41% | 0.17% |
| Proliferative | 1.18% | 1.40% | 1.58% | 1.26% | 1.73% | 4.51% |
| myCAF | 1.16% | 0.17% | 3.94% | 5.14% | 0.71% | 0.03% |
| Acinar | 0.68% | 0.00% | 3.92% | 0.11% | 0.00% | 0.38% |

**Supplementary Table S3.** Percentage of immune cells in mouse KRAS G12C and G12D tumors tissues.

| Annotation | G12C_6304 | G12C_7825 | G12C_9726 | G12D_4956 | G12D_57614 | GSM3577885 |
| --- | --- | --- | --- | --- | --- | --- |
| B | 4.73% | 3.81% | 36.30% | 8.11% | 14.55% | 1.26% |
| DC | 9.03% | 3.88% | 3.21% | 21.08% | 3.94% | 28.46% |
| Macrophage | 2.09% | 1.12% | 1.07% | 1.08% | 2.12% | 36.83% |
| Monocyte | 5.98% | 5.99% | 2.80% | 21.62% | 8.79% | 19.45% |
| Neutrophil | 71.06% | 78.28% | 20.00% | 44.32% | 60.61% | 11.34% |
| Plasma | 0.67% | 0.25% | 2.14% | 0% | 3.03% | 0.35% |
| T | 6.44% | 6.68% | 34.49% | 3.78% | 6.97% | 2.32% |

## Notes

### Summary of Updates

Figure 2, 3,4 and supplementary Figure S4 updated; Authors added.

